# Rhs-toxin Abundance, Diversity, and Function in Four Genera of Plant Pathogenic Bacteria: *Xanthomonas, Ralstonia, Pectobacterium,* and *Dickeya*

**DOI:** 10.1101/2025.05.08.652813

**Authors:** Andrea Gómez Cabrera, Yaken Obaydeh Ameen, Katherine D’Amico-Willman, Prasanna Joglekar, Jose Huguet-Tapia, Ignazio Carbone, Alejandra I. Huerta

**Affiliations:** Department of Entomology and Plant Pathology, NC State University, Raleigh, NC 27695, USA; Department of Plant Pathology, University of Florida, Gainesville, FL 32611, USA; Center for Integrated Fungal Research, Department of Entomology and Plant Pathology, North Carolina State University, Raleigh, NC 27695, USA

## Abstract

Rearrangement hotspot (Rhs) toxins are polymorphic bacterial toxins that inhibit competitor microbes, playing a critical role in interbacterial competition. Each toxin comprises a conserved N-terminal region for translocation and a hypervariable C-terminal domain harboring toxic activity, with a cognate immunity gene positioned downstream to prevent self-killing. The ubiquity and variability of Rhs toxins across bacterial genera suggest they contribute significantly to microbial fitness and niche exclusion. However, their distribution, diversity, and functional roles remain poorly understood in plant-pathogenic bacteria. Here, we used a profile Hidden Markov Model to systematically mine genomes from four agriculturally important plant-pathogenic genera: *Xanthomonas*, *Ralstonia*, *Pectobacterium*, and *Dickeya*, identifying 604, 294, 255, and 113 Rhs homologs, respectively, across 343 genomes. N-terminal sequence classification revealed multiple distinct families, including lineage-specific groups exclusive to *Xanthomonas* and *Ralstonia*. Notably, these were linked to the type II secretion system, diverging from the canonical association with type VI secretion. C-terminal domain analyses via sequence similarity networks revealed both conserved and lineage-specific toxic variants. *Xanthomonas* strains encoded the most diverse repertoire, including predicted DNases, RNases, proteases, and deaminases. However, the functions of 69.6% of C-terminal domains remain uncharacterized. Contrary to our initial hypothesis that soilborne bacterial pathogens would encode more abundant and diverse Rhs toxins due to intense microbial competition in soil, foliar pathogens exhibited greater Rhs diversity. This suggests that aboveground plant environments may impose stronger selective pressures for Rhs toxin diversification. These findings highlight the unexplored potential of Rhs toxins in shaping microbial ecology and underscore the need for functional characterization to elucidate their roles in bacteria-microbiome interactions.

**Author Summary:** Bacteria constantly interact and compete for resources with other microbial agents in plant-associated microbiomes. One way they can enhance their competitive fitness is by producing proteinaceous toxins that can harm or kill rival cells. One group of toxins used for intraspecies competition is the Rhs toxins, which have a unique structure and diversity of enzymatic actions encoded in their hypervariable protein tip. To better understand the diversity and abundance of Rhs toxins in plant pathogenic bacteria, a computational pipeline was developed and used to analyze publicly available genomes from four major bacterial plant pathogenic genera. Results confirm the ubiquity of Rhs toxins in bacterial genomes and show the lack of Rhs toxins in genomes from unique species. Furthermore, some Rhs toxin enzymatic functions were unique to a particular genus. The data presented here suggest that some Rhs toxins may be secreted through alternative pathways beyond the well-known Type VI secretion system. Together, this study on the abundance and diversity of Rhs toxins in plant pathogenic bacterial genera highlights the complexity and predicted functional diversity of Rhs toxins and provides fundamental knowledge to test hypotheses on the role Rhs toxins play in microbial ecology, community structure, and evolution in the context of plant disease.

## Introduction

Microbial community composition is shaped by the relationships and interactions between and among constituent members of that community. These dynamic interactions ranging from beneficial to mutualistic to antagonistic, negatively and positively impact a couple, a few, or all community members in microbe-rich environments [1]. These stringent bacteria-microbiome interactions place evolutionary pressures on individual bacterial cells and species to evolve diverse competitive mechanisms to enhance their fitness [2]. This includes the development of an arsenal of antimicrobial compounds that disrupt competitor cell physiology. These include antibiotics, bacteriocins, and the more recently described polymorphic toxins, including the Rhs toxins [3]. The decreased effectiveness of antibiotics and copper-based agrochemicals in treating human and plant bacterial infections make Rhs toxins and other polymorphic toxins an attractive alternative for developing targeted management compounds [4–6]. However, before these compounds are developed into disease management tools, fundamental knowledge of their ecological significance is essential, including their abundance, diversity, function, and host range within and across bacterial species and genera.

The typical Rhs locus (Rhs system) in gram-negative bacteria is comprised of three genes: *rhs, immunity,* and *chaperone*. The *rhs* gene encodes for an approximately 1,500 amino acid (aa) long polymorphic toxin composed of a conserved GC-rich N-terminal region that facilitates toxin translocation from the producing cell to the target cell [7–9]. The tyrosine/aspartate (YD) peptide repeats encoded at the N-terminus define the Rhs toxin family [2, 10]. In gram-negative bacteria, a conserved 10 amino acid peptide motif, PxxxxDPxGL, sharply demarcates the N-terminal domain from the GC-poor Rhs C-terminal hypervariable tip (Rhs-CT) [2]. The C-terminal tip encodes for the active killing domain of the Rhs toxin. The *rhs* gene is followed by an *immunity* gene that encodes for a cognate immunity protein that protects the producing cell from autoinhibition [2, 11, 12]. Together, these two elements make a toxin anti-toxin system, which, together with the *chaperone* gene completes the Rhs toxin system [13].

Occasionally, one or more orphan modules can be found directly downstream of the full-length Rhs toxin and cognate immunity gene pair. Orphan modules do not contain the full-length N-terminal. They do, however, contain homology to the 3’ extremity of the N-terminal, including the conserved 10 aa motif, and encode alternative toxic domains in their Rhs-CT and associated immunity protein [2]. In-vitro, the C-terminal tip of the full-length Rhs toxin can be displaced via homologous recombination with an orphan module to generate new functional toxins [14]. This phenomenon suggests that, in nature, Rhs toxins are undergoing homologous recombination, thus contributing to Rhs toxin diversification and bacterial fitness [2].

Rhs toxins can be translocated from the cytoplasm of the producing cell directly into a prokaryotic cell via the type VI secretion system in a contact-dependent manner with the help of chaperone proteins, like VgrG [15, 16]. The T6SS is composed of 13-14 core proteins (TssA-TssM, PAAR), that form a puncturing apparatus, a cytoplasmic contractile sheath, a cytoplasmic baseplate, and a membrane complex that together create a membrane-embedded, spear-like weapon that translocates the Rhs toxin [17]. The T6SS-associated *vgrG*, *hcp*, and *paa*r genes are essential for Rhs toxin translocation and are often found contiguous to Rhs toxin systems [16, 18–20].

Functionally characterized Rhs-CT tip enzymatic actions include DNases, RNases, glycosylases, and peptidases [10]. For example, one Rhs-CT tip in *Escherichia coli* strain EC869 encodes a DNase that kills *E. coli* strain MC4100 through DNA degradation [21]. An Rhs toxin in S*almonella enterica* inhibits protein synthesis in an *E. coli* strain through ADP-ribosylation, thus altering the activity and function of proteins in the target cell [22]. Bioinformatic pipelines designed by Zhang et al. [10] and Danov et al. [23] predict nucleases are the most abundant and widespread mode of action among Rhs-toxins [10, 24]. However, many Rhs-CT tips remain uncharacterized.

Studies describing the roles and molecular mechanisms of Rhs toxins are focused primarily on translocation via the T6SS and most often on individual human and animal pathogens as models [25–29]. Very few provide insight into Rhs toxin abundance, diversity, and function in agriculturally important bacterial strains, species, and genera at the population level. Here, we aimed to elucidate Rhs toxin abundance, diversity, and predicted function in four genera of economically important bacterial plant pathogens. To accomplish this goal, we designed and optimized a bioinformatic pipeline that accelerated genome mining for Rhs toxins from 509 genomes from 21 plant pathogenic *Xanthomonas* species, 133 genomes from 5 *Ralstonia* species, 95 genomes from 15 *Pectobacterium* species, and 61 genomes from 11 *Dickeya* species.

## Results and Discussion

### The genomes of plant pathogenic bacteria encode an array of full-length Rhs toxins and numerous orphan Rhs-CT tips

A total of 509 *Xanthomonas*, 133 *Ralstonia*, 95 *Pectobacterium*, and 61 *Dickeya* closed genomes were initially downloaded from NCBI (Table S1). Following an average nucleotide analysis to remove clonal genomes (100% similarity) and additional manual curation as described in the methods section, a total of 165, 87, 60, and 31 complete single chromosome genomes were retained for downstream analysis, respectively (Fig 1A; Table S1 and S2). Rhs toxins were not recovered from genomes in *X. phaseoli, P. punjabense, D. aquatica*, *D. lacustris*, *D. poaceiphila*, *R. pickettii, R. mannitolilytica,* and *R. insidiosa* using the methods described here and were removed from further analysis (Table S2). Of the final 343 genomes analyzed, 131 encoded at most two Rhs toxins, 183 encoded three to seven, and 28 encoded more than eight (Table S2). Genomes of *P. parmentieri* encoded the highest number of Rhs toxin sequences with 13 in strains QK-5 and WC19161, followed by *R. solanacearum* strain CFBP2956 with 12, and *X. citri* strain UnB-XtecTG02-2 and *X. oryzae* strains BB156-2 and BB151-3, with 11 (Fig 1B; Table S2).

**Fig 1.**
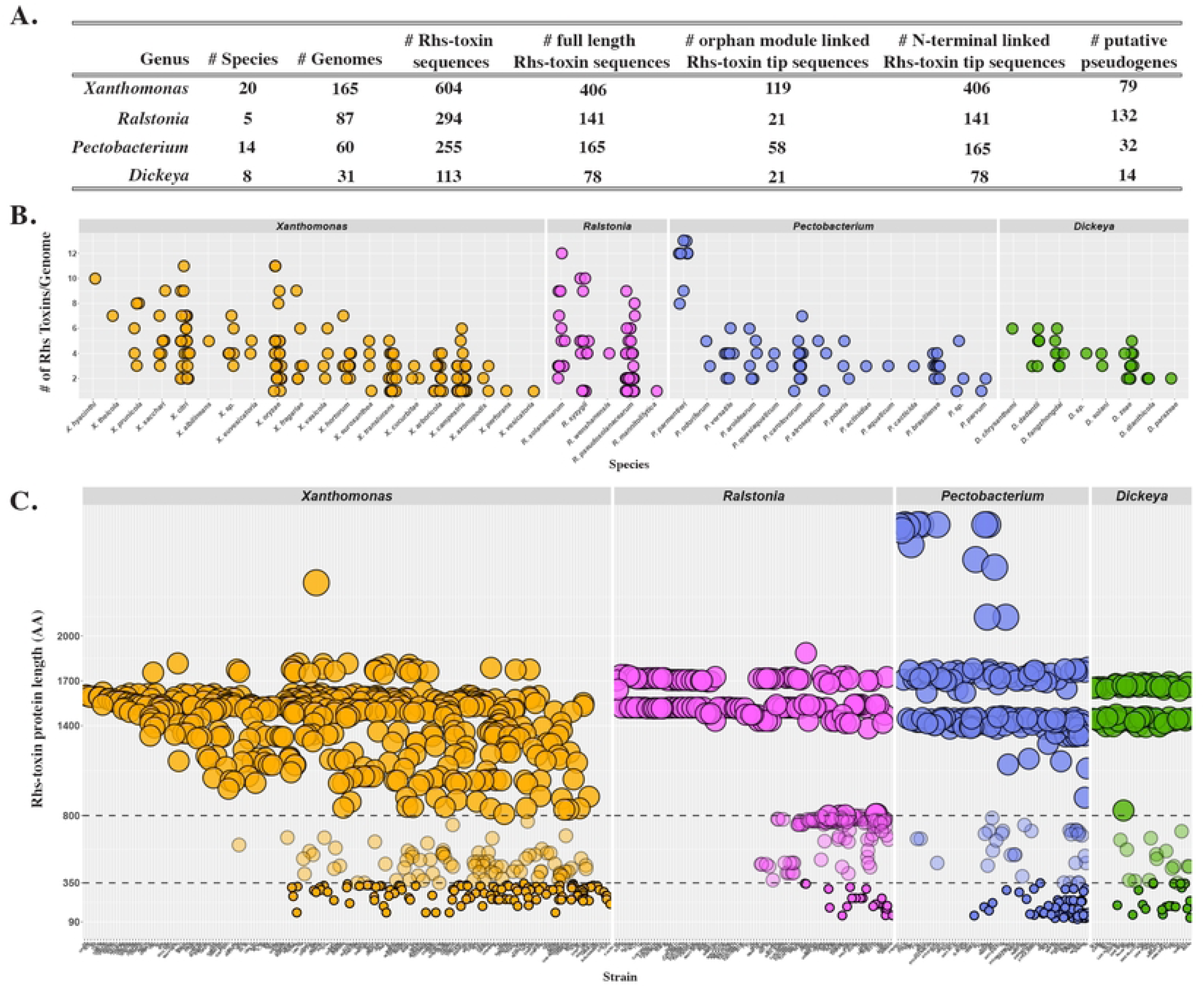
Rhs toxin sequence abundance and size categories in four genera of plant pathogenic bacteria: *Xanthomonas, Ralstonia*, *Pectobacterium,* and *Dickeya*. **(A)** Summary table with number of species, genomes, and Rhs toxin sequence categories post data curation. (B) Number of Rhs toxin sequences per species for each genus in this study. On the x-axis are species, and on the y-axis is the number of Rhs toxin sequences per genome. **(C)** Rhs toxin protein lengths within, between, and among bacterial genomes for each of the four genera. On the x-axis, individual genomes and on the y-axis, the protein length in amino acids (aa) for each Rhs toxin; each dot represents a single Rhs-toxin sequence. The size of a dot correlates to Rhs toxin category (e.g., protein length): <350 aa (small dots) represent Rhs toxin orphan module C-terminal tips; 350 ≤ 799 aa (medium dots) represent Rhs toxin pseudogenes; and > 800 aa (large dots) represent full-length Rhs toxins. The dashed lines intercepting the y-axis at 350 aa and 800 aa separate the orphan module C-terminal tips, Rhs-toxin pseudogenes, and full-length Rhs-toxin sequences. Each dot is color-coded based on genus.

### Rhs toxin and orphan module abundance are highest in *Xanthomonas*

To better understand the abundance of full-length Rhs toxin and orphan modules in these four genera of plant pathogenic bacteria, protein sequences were separated into three categories based on length: Full-length Rhs toxins (>800 aa), pseudogenes (360-799 aa), and orphan modules. (≤350 aa) as described by Steele et al. [30] (Fig 1A and 1C; Table S3). Among the 604 Rhs toxin sequences identified across 20 *Xanthomonas* species, 406 were full-length Rhs toxins, 119 were orphan modules, and 79 were pseudogenes. Rhs-toxin abundance and diversity were the highest among *Xanthomonas* compared to soilborne *Ralstonia, Pectobacterium,* and *Dickeya*.

*Xanthomonas* also had the highest number of predicted orphan modules, suggesting more competition and possible Rhs CT tip recombination among species within the genus (Fig 1C). This raises the hypothesis that foliar plant pathogens experience more bacterial competition, driving Rhs toxin diversity. While the abundance of Rhs toxins observed in *Xanthomonas* supports an intriguing hypothesis about ecological pressures in foliar pathogens, this interpretation should be cautiously approached. The underlying dataset is skewed by the overrepresentation of Xanthomonas genomes in public databases, which may artificially inflate diversity and abundance metrics. Future studies should employ normalization strategies or balanced sampling across genera to rigorously test this hypothesis.

About 57% of all *Xanthomonas* genomes encoded at least one full-length Rhs toxin but no orphan modules (Fig 1C; Table S3). The non-orphan module encoding genomes predominated among strains in *X. cucurbitae, X. euroxanthea, X. translucens,* and *X. vesicatoria* species that are pathogenic on cucurbit, walnut, wheat, and pepper and tomato, respectively. Interestingly, most of these species are known to infect only one host except for *X. vesicatoria,* which infects pepper and tomato, and all are predicted to be seed-transmitted [31, 32]. The lack of orphan modules and the small repertoire of full-length Rhs toxins suggest these species undergo little intraspecies competition or rely on alternative competition mechanisms.

The genome with the most orphan modules in *Xanthomonas* was *X. hyacinthi* strain CFBP 1156, the causal agent of yellow disease in *Hyacinthus,* with eight orphan modules and two full-length Rhs toxins. The abundance of orphan modules in this strain suggests it is under greater competitive pressures and thus may have a high competitive fitness; however, little is known about the biology and genetics of this pathogen to make any conclusion as to why it harbors this large number of orphan modules [33–36].

### Rhs toxin pseudogenes are most abundant in *Ralstonia*

Among the 294 Rhs toxin sequences identified across the six *Ralstonia* species analyzed, 141 were full-length Rhs toxins, 21 were orphan modules, and 132 were pseudogenes (Fig 1A and C; Table S3). Rhs toxin pseudogenes are truncated at the N-terminal region, possibly preventing their transcription and activity [30]. *Ralstonia* exhibited the highest number of pseudogenes among the four genera analyzed. This indicates that its strains may experience greater intraspecies competition, leading to stronger negative selection, forcing strains in the genus to diversify their Rhs toxin repertoire more often and acquire new Rhs toxins while inactivating others. This finding is consistent with the well-known genomic plasticity of the *Ralstonia solanacearum* species complex (RSSC), which contributes to the pathogens’ broad host range and remarkable environmental adaptability [37–39].

It remains unclear how Rhs toxin abundance impacts a strain’s competitive fitness. Interestingly, *R. solanacearum* strain K60^T^, the type strain for the species, encoded nine Rhs-toxins: three full-length Rhs toxins, one orphan module, and five pseudogenes (Table S3). GMI1000 encoded six Rhs toxin sequences: two full-length Rhs toxins, and four pseudogenes, while UW551 encoded only one full-length Rhs toxin and two putative pseudogenes. Interestingly, K60 was found to directly inhibit the growth of both GMI1000 and UW551 in overlay inhibition assays and outcompeted both strains in tomato rhizospheres and stems [40]. Echandi et al [41] also confirmed K60’s competitive fitness against other strains in the RSSC. These findings raise the hypothesis that Rhs toxin abundance plays a role in K60’s competitive advantage against closely related strains.

### *Pectobacterium* strains encode unusually long Rhs toxins

The Soft Rot Pectobacteriaceae, *Pectobacterium,* and *Dickeya*, encoded 165 and 78 full-length Rhs toxins, 58 and 21 orphan modules, and 32 and 14 pseudogenes, respectively (Fig 1A and 1C; Table S3). Among the four genera investigated, the number of genomes available for *Dickeya* was the lowest, possibly confounding the small number of complete Rhs-toxins, pseudogenes, and orphan modules found for this genus (Table S1 and S2). Twelve *g*enomes from among *P. carotovorum*, *P. brasiliense*, *P. odoriferum*, *P. parvum*, and *P. versatile,* and one from a *Pectobacterium* with no species designation encode unusually long (> 2000 aa) Rhs toxin sequences. These were similar in length to one Rhs toxin sequence found in the genome of *Xanthomonas* strain AM6 (Fig 1C; Table S3). The 500 amino acid extension of *Xanthomonas* Rhs-toxin OCJ37_RS10125 in strain AM6 encode SpvB, TcdB, and TcdC domains, all associated with known bacterial toxins [9, 42–44]. An NCBI BLASTp search using the unusually long Rhs toxin sequences identified similarly long Rhs toxins in *Serratia marcescens*, *S. bockelmannii*, *Photobacterium alginolyticum*, and *Denitrovibrio acetiphilus* [11, 45–47].

The longer N-terminal region in these sequences suggests that these Rhs toxins may use a secretion mechanism independent of the T6SS typically predicted for most Rhs toxins [48]. Similar toxins in *Salmonella enterica* are suspected to be translocated via the type III secretion system [49]. Similarly, Steele et al. [25] noted the presence of these long Rhs toxins in the genomes of *Apibacter* spp., but they predict these Rhs toxins are translocated via the type IX secretion system [25].

### Rhs toxin families correlate to Rhs toxin system loci, synteny, and predicted secretion mechanism

To investigate the translocation of Rhs toxins in each of the four genera of plant pathogenic bacteria in this study, we first classified Rhs toxins into families using the N-terminal “core region” defined by the two peptide motifs RVxxxxxxxG and PxxxxDPxGL [2, 25]. For each genus, two distinct methods (IQ-TREE and RAxML) were used to infer and confirm Rhs toxin clade phylogenies based on N-terminal core region alignment (Fig S1-S4). In IQ-TREE clades with bootstrap values higher than 95% and phylogenetic distance of 0.798 were considered well supported and were used to define Rhs toxin families for all four genera (Fig 2A, 2C, 2D, and 2E; Fig S5-S8; Table S4). This analysis was repeated using RAxML with a general time-reversible (GTR) matrix-based model of amino acid substitution rates and 1,000 bootstrap replicates [50]. All Rhs family clades were well supported with bootstrap values higher than >90% and phylogenetic distances of 2.28, 16.4, 1.41, and 12.0 for *Xanthomonas, Ralstonia, Pectobacterium,* and *Dickeya*, respectively (Fig S1B, S2B, S3B, and S4B). With these phylogenetic analysis parameters, no differences were observed in family clades designation, supporting defined Rhs toxin families for the four genera.

**Fig 2.**
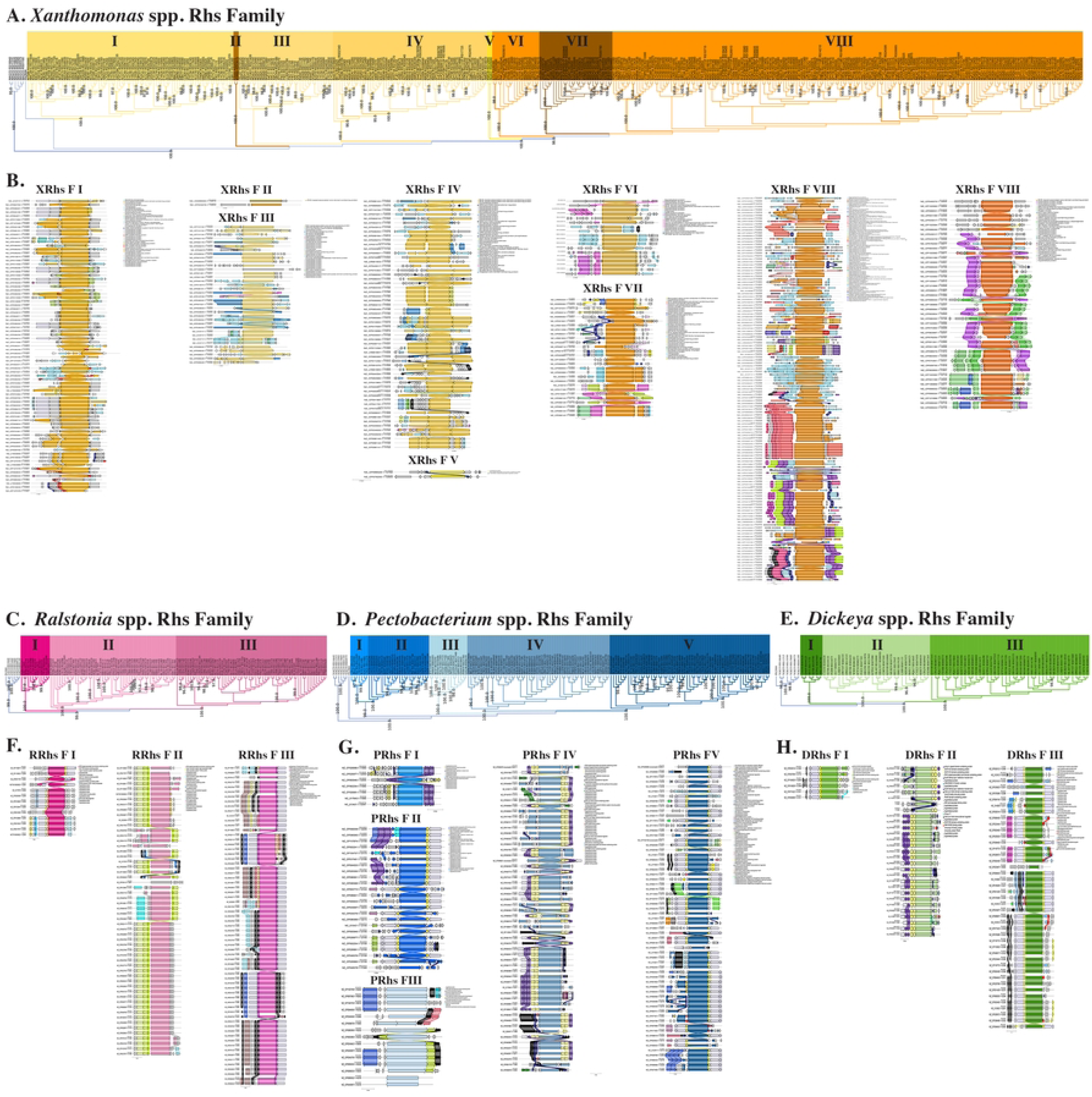
Classification of Rhs toxin families in (A) *Xanthomonas*, (C) *Ralstonia*, (D) *Pectobacterium*, and (E) *Dickeya*. Rhs toxin clade phylogenies were based on N-terminal core region alignment of 406, 141, 165, and 78 protein sequences for each of the four genera, respectively, in addition to seven Rhs toxin sequences from *E. coli* and *P. aeruginosa*, used an an outgroup. All sequences were aligned using MAFTT and phylogenies inferred using IQ-TREE for each genus. Each defined Rhs family is indicated by roman numerals within the corresponding phylogenetic and the outgroup clade is represented in white. All clade families are supported by bootstrap values higher than 95% and phylogenetic distances of 0.798. Gene synteny among Rhs toxin loci for the eight, three, five, and three Rhs toxin families identified for (B) *Xanthomonas,* (F) *Ralstonia*, (G) *Pectobacterium*, and (H) *Dickeya*. A locus defined as 5000 bps upstream and 3000 bps downstream of each Rhs toxin sequence within each family was extracted from the respective genomes. All Rhs toxins and associated up- and downstream sequences were aligned and visualized using clinker (Gilchrist and Chooi 2021). Each Rhs toxin family within a genus is represented by a different shade of orange, pink, blue, or green.

In *Xanthomonas,* eight distinct Rhs families were identified using the best fit model Q.pfam+F+G4 selected by IQTree, named *Xanthomonas* Rhs Family (XRhs F) I-VIII (Fig 2A; Fig S1A and S5; Table S4). In *Ralstonia*, three distinct Rhs families were identified using the best fit model JTT+I+G4 by IQTree, named *Ralstonia* Rhs Family (RRhs F) I-III (Fig 2C; Fig S2A and S6; Table S4). In *Pectobacterium,* five distinct Rhs families were identified using the best fit model Q.insect+F+G4 by IQTree, named *Pectobacterium* Rhs Family (PRhs F) I-V (Fig 2D; Fig S3A and S7; Table S4). In *Dickeya*, three distinct families were identified using the best fit model Q.plant+F+I+G4 by IQTree, named *Dickeya* Rhs Family (DRhs F) I-III (Fig 2E; Fig S4A and S8; Table S4).

*Xanthomonas* XRhs F VIII, composed of 180 Rhs toxin sequences from 118 *Xanthomonas* genomes and 18 species, was the most prominent Rhs toxin family across all genera in this analysis (Fig 2B; Fig S16; Table S4). Toxins in this family were associated with the *gspG* gene, which encodes the periplasmic pseudopilus of the Type II secretion system (T2SS) [51]. Thus, we predict that toxins in this family are translocated via the T2SS, which is responsible for delivering toxins and other enzymes to the bacterial cell surface or extracellular space in gram-negative bacteria [52]. This contrasts with the contact-dependent mechanism of the T6SS. Similarly, all the 56 Rhs toxins in *Ralstonia* RRhs F II were also linked to the *gspG* gene (Fig 2F; Fig S18; Table S4). Although Rhs toxin secretion via the T2SS has not been functionally confirmed, indirect evidence by Huerta et al. [40] and Marutani-Hert et al. [53] supports this finding. In both studies, the authors showed that cell-free supernatant from bacterial overnights inhibits the growth of target strains via an overlay inhibition assay. The activity was confirmed to be mediated via Rhs toxins, bypassing the contact-dependent nature of the T6SS [40, 53].

Functional characterization of these toxins within the context of the T2SS and T6SS mutants will confirm these observations. Nonetheless, the data presented here support the hypothesis that Rhs toxins have secretion and delivery mechanisms other than the T6SS [25]. Not surprisingly, all remaining Rhs toxin families, regardless of species and genus, were linked and showed different degrees of synteny to T6SS-associated genes, supporting a contact-dependent translocation mechanism for toxins in those families (Fig 2B, 2F, 2G, and 2H; Fig S9-S15, S17, and S19-S27; Table S4).

None of the Rhs toxin families in *Pectobacterium* or *Dickeya* were linked to genes associated with the T2SS (Fig 2G and 2H; Fig S20-S27). This raises the question, why do *Pectobacterium* or *Dickeya* not encode T2SS translocated Rhs Toxins? These two genera belong to the Soft Rot Pectobacteriaceae, which rely on the T2SS for the secretion of essential virulence factors—cell-wall degrading enzymes, primarily pectinases—that break down pectin in the middle lamella of plants, leading to tissue maceration and soft rot [52, 54]. This observation indicates that cell wall-degrading enzymes hold greater evolutionary significance for soft rotting *Pectobacterium* and *Dickeya* than the diversification of Rhs toxin translocation mechanisms. All the Rhs toxins in *Pectobacterium* and *Dickeya* were physically co-localized with two genes of the T6SS, *vgrG* and *hcp* (Fig 2G and 2H). Both *vgrG* and *hcp* co-localize to Rhs toxin loci and were often found upstream of the N-terminal of XRhs F I; RRhs F I and III; PRhs F I, II, IV, and V; and DRhs F I- III (Fig 2B, 2F, 2G and 2H; Fig S9, S17, S19-S21, and S23-S27).

### Domains of Unknown Function (DUF) determine Rhs toxin family phylogenetic structure

Four genes encoding Domains of Unknown Function (DUF): DUF1795, DUF2169, DUF4150, and DUF4123, were prominent among Rhs toxin families across the four bacterial genera [55]. In these families, a DUF protein preceded the Rhs toxin. In the Rhs toxins phylogenetic analysis the toxins with the same DUF protein clustered together. This observation suggests that DUF proteins drive Rhs toxin family phylogenetic structure. In Rhs toxin systems, DUF proteins have been proposed to aid in the recruitment of Rhs toxins to the T6SS apparatus and, thus, function as Rhs toxin chaperones [56]. Among these are DUF1795 proteins, which in this analysis was associated with RRhs F I, PRhs F II and V, and DRhs F I and III (Fig 2F, 2G and 2H; Fig S17, S21, S24-S25, and S27). The DUF1795 proteins, recently classified as Eag proteins, are predicted chaperone proteins that bind and stabilize the N-terminal of the Rhs toxins and facilitate their interactions with VgrG [47, 57, 58]. Similarly, DUF4123 proteins were associated with two unique Rhs toxin families: *Pectobacterium* PRhs F IV and *Dickeya* DRhs F II (Fig 2G and 2H; Fig S23 and S26). DUF1795 and DUF4123 proteins have been proposed as chaperones of T6SS secreted effectors [59]. However, structural and functional studies are necessary to elucidate how these chaperones interact with Rhs toxins and the different molecular components of the T6SS. Lastly, DUF4150 and DUF2169 proteins are also predicted to be Rhs toxin chaperones associated with Ankyrin proteins in PRhs F I and IV and in DRhs F II (Fig 2G and 2H; Fig S20, S23 and S26). Ankyrin proteins have been found to mimic antibodies, are highly stable, and can withstand degradation by proteases and harsh chemicals [10, 60, 61]. Perhaps these Ankyrin-associated chaperone proteins protect the Rhs toxin from degradation inside the producing cell and the cytoplasm of the target cell.

*Pectobacterium* PRhs F III housed the unusually long N-terminal Rhs toxins (>2,000 aa) that are encoded in the genomes of *P. versatile*, *P. odoriferum*, *P. caratovorum*, *P. parvum*, and *P. brasiliense* (Fig 1C and 2G; Fig S22; Table S3 and S4). This family was not associated with any genes related to a known secretion system. However, they were associated with a ssDNA-binding protein (SSB1): an Hsp70-type chaperone protein shown to assist in the folding and assembly of newly synthesized proteins in the bacterial cytosol [62]. This suggests these longer Rhs toxins require additional proofreading and quality control compared to the 1,700 aa Rhs toxins.

Transposable element insertion in genes that code Rhs toxins, orphan modules, or their chaperones can impact their expression and function [63, 64]. Thirteen families had Rhs toxins associated with transposable elements, including XRhs F I, XRhs F III, XRhs F IV, XRhs F VI- XRhs F XIII, RRhs F I-RRhs F III, PRhs F V, DRhs F I, and DRhs F III (Fig 2B, 2F, 2G, and 2H; Fig S9, S11-S12, S14-S19, S24-S25, and S27). The most prominent transposable elements belonged to IS3, IS5, IS30, and IS1595, each with distinct characteristics that can impact gene mobility, regulation, and expression. As Lerat et al. (2005) noted, most genes disrupted by transposable elements tend to become pseudogenes [65]. This is evident among *rhs* genes, as we show here and as reported by Steele et al. (2017; 2021).

### Homology and predicted enzymatic function of Rhs toxin C-terminal tips across strains, species, and genera

The Rhs-CT tips feature the toxic domain responsible for disrupting the biological processes of the target cell [66, 67]. However, little is known about the intra- and inter-species distribution of Rhs CT tips, which can ultimately impact competitive outcomes among strains, particularly if the target cell carries a cognate *immunity* gene. To better understand Rhs-CT tip distribution among strains in a genus and to predict toxic functions, we performed an all-vs-all blastp analysis followed by a similarity network analysis that grouped Rhs-CT tips into clusters based on >85% similarity. A total of 183 unique Rhs-CT tips, named XRhs1-Xrhs183, were identified among the 604 Rhs-CT tip sequences retrieved from *Xanthomonas* genomes. Of those, 86 Rhs-CT tips were singletons present in only one strain of the genus Xrhs98-Xrhs183 (Fig 3A and 4A; Table S3). The remaining *Xanthomonas* Rhs-CT tips were present in at least two strains. Specifically, 26 Rhs-CT tips were present in two *Xanthomonas* strains, 54 Rhs-CT tips were shared among three to seven strains, and 18 were present in more than eight strains (Xrhs1- Xrhs18) (Fig 3A; Table S3).

**Fig 3.**
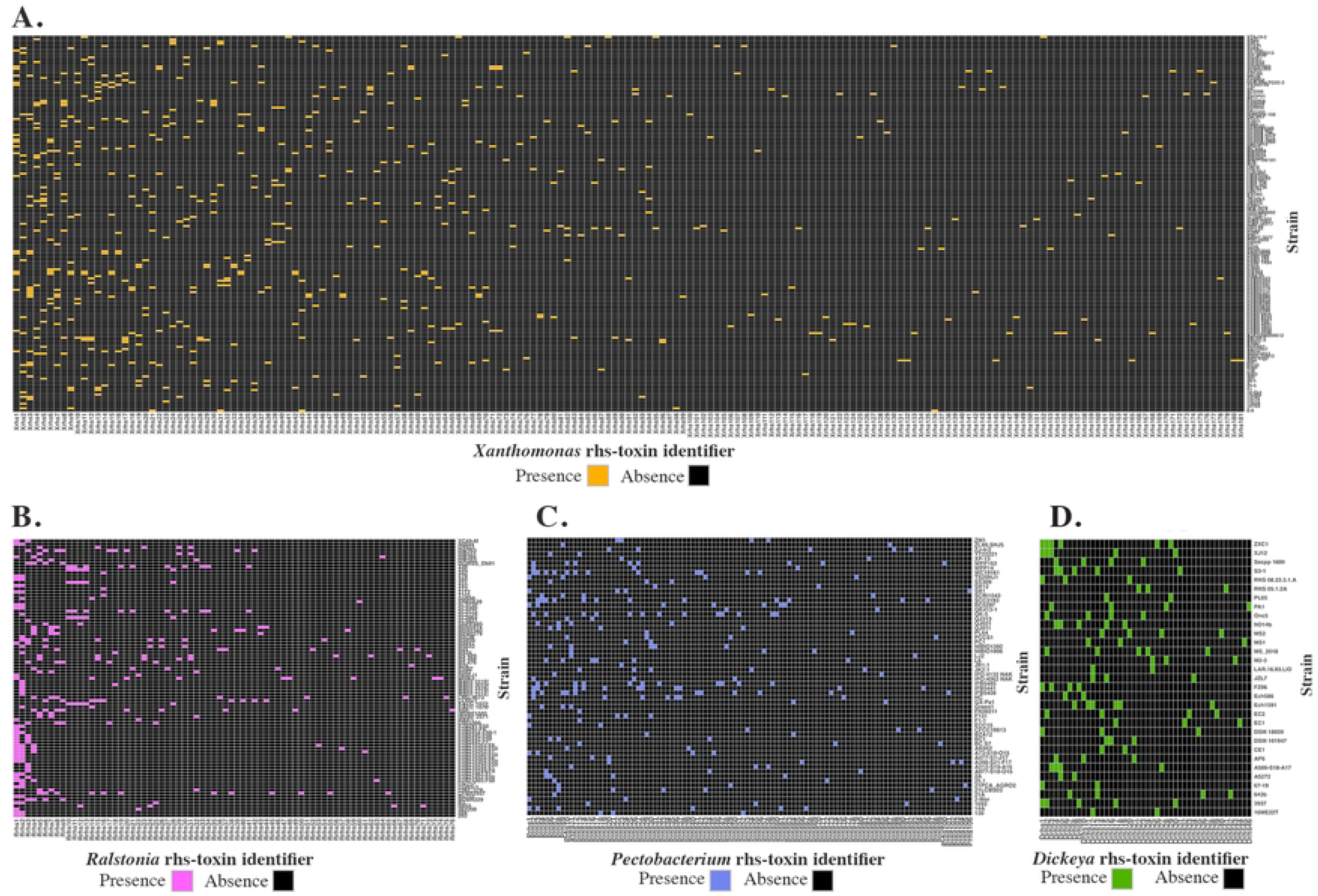
Presence and absence plots of Rhs-CT tips across strains of (A) *Xanthomonas*; (B) *Ralstonia*; (C) *Pectobacterium*; and (D) *Dickeya*. To determine protein homology a blastP analysis was used on 604, 294, 255, and 113 C-terminal tip sequences from 165, 87, 60, and 31 genomes from each genus, respectively. Rhs-CT tips were considered homologous if they shared more than 85% amino acid similarity. These were grouped into Rhs toxin tip identifiers as follows: Xrhs1–Xrhs183 for *Xanthomonas*, Rrhs1–Rrhs76 for *Ralstonia*, Prhs1–Prhs106 for *Pectobacterium*, and Drhs1–Drhs46 for *Dickeya*. On the x-axis, the Rhs toxin tip identifier and the y-axis strains used in the analysis. A solid colored box indicates the presence of an Rhs-CT tip in a given strain: orange for *Xanthomonas*, pink for *Ralstonia*, blue for *Pectobacterium*, and green for *Dickeya*. A black box indicates the absence of that Rhs-CT tip in the corresponding strain.

**Fig 4.**
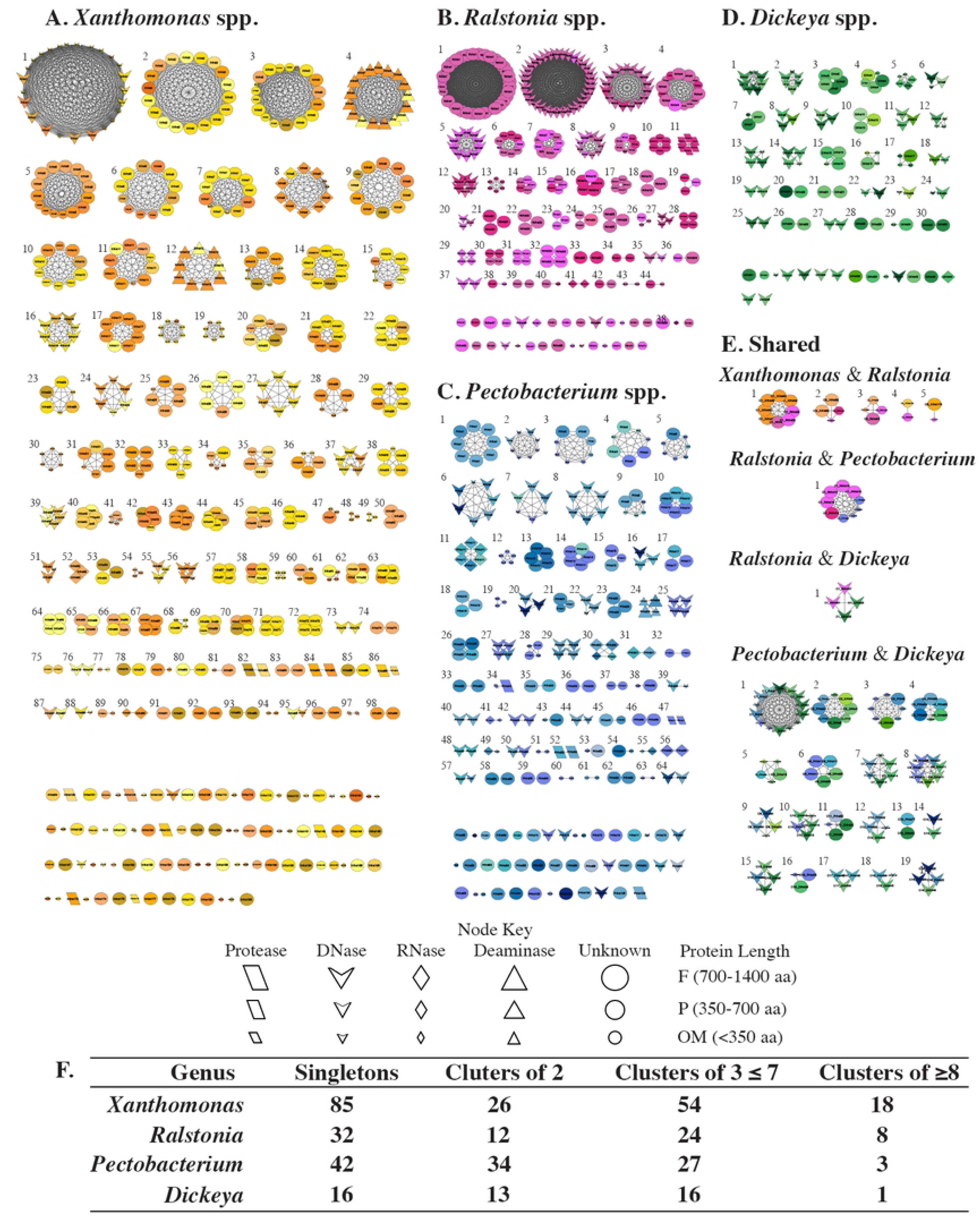
Homology and predicted enzymatic function of Rhs toxin C-terminal tips in (A) *Xanthomonas*, (B) *Ralstonia*, (C) *Pectobacterium*, and (D) *Dickeya*. An all vs all BlastP analysis was used to perform a similarity network analysis for 604, 294, 255, 113 Rhs-CT tips among and (E) between genera. Rhs-CT tip identifier was overlaid on each node in accordance with Fig 3. Node color, shape, and size represent a bacterial species, the predicted enzymatic function, and the protein length for each Rhs-CT tip, respectively. (F) Summary table of Rhs CT tip clusters for each genus.

The most abundant Rhs-CT tip in the genus *Xanthomonas* was Xrhs1, which was present in 25 different *Xanthomonas* strains. This included 12 *X. citri* (XcmN1003, XcmH1005, TX160149, NT17, MSCT, LMG7439, DAR73886, DAR72029, CFBP7112, CFBP 2036, AW15), three *X. vasicola* (Xv1601, NCPPB 2649, NCPPB 1060), three *X. prunicola* (MAI5037, CIX383, CIX249), one *X. campestris* (NCPPB 4379), and one *Xanthomonas* with no species designation (WG16) (Fig. 3A; Table S3). The Xrhs1 tips are predicted DNases encoded by a Tox-REase-7 domain [10, 68] (Fig4A; Fig S28; Table S3). The second (Xrhs2) and third (Xrhs3) most abundant Rhs-CT tips were present in 18 strains (Fig. 3A; Table S3). The Xrhs2 tip, with unknown function, was encoded by two *X. arboricola* (YchA and 1314c), four *X. translucens* (Xtu-UPB513, XtKm33, MAI5034, ICMP 16317), one *X. hortorum* (108), seven *X. campestris* (M28, GBBC 3077, CFBP 8444, 85-1016-10, 12049, 576), two *X. euroxanthea* (CPBF 424, CPBF 367), one *X. euvesicatoria* (LMG930), and one *X. hyacinthi* (CFBP 1156). Similarly, the Xrhs3 tips have unknown function (Fig 4A; Fig S28; Table S3). However, Xrhs3 tips encode the DUF4329, which has been linked with two conserved protein domains pfam05593 and pfam03527 associated with Rhs repeats in the N-Terminal of the Rhs toxin [29, 69] (Table S3). Unlike Xrhs2 tips which were found across seven species of *Xanthomonas*, Xrhs3 tips were encoded in four species: *X. hortorum* (108, jj2001)*, X. citri* (DAR73886, DAR72029, CFBP7119, CFBP7112, CFBP7111, CFBP6991, CFBP6990, AW15, 8ra)*, X. campestris* (85-10), *X. arboricola* (1314c, 1311a, 305) and a *Xanthomonas* with no species designation (WG16) (Table S3). Unsurprisingly, the function of 143 Rhs-CT tips remains to be elucidated, raising the questions: *what is the enzymatic activity of these toxins* and *why are some conserved across different species*?

A total of 76 unique Rhs-CT tips, named Rrhs1-Rrhs76, were identified among the 294 Rhs-CT tip sequences retrieved from the 87 *Ralstonia* genomes analyzed. Of those, 32 Rhs-CT tips were singletons and present in only one strain of this genus (Rrhs44-Rrhs76) (Fig 3B; Table S3). The remaining *Ralstonia* Rhs-CT tips were present in at least two strains. Specifically, 12 RRhs tips were present in two *Ralstoina* strains, 24 Rrhs tips were shared among three to seven strains, and eight tips were present in eight or more strains (Rrhs1-Rrhs8) (Fig 3B; Table S3).

The most abundant Rhs-CT tip among strains in the RSSC was Rrhs1, which was present in 36 strains (Fig 3B; Table S3). The tips that comprise Rrhs1 have an unknown function, but their presence across multiple strains suggests it is vital for this pathogen’s fitness (Fig 4B; Fig S29; Table S3. All but one of the Rhs-CT tips in Rrhs1 are present in *R. pseudosolanacearum* strains. These were isolated from tomato, tobacco, peanut, potato, and infected soils across different geographic regions, including Nigeria, China, Korea, and the Philippines (Table S1 and S3). The 36th Rhs-CT tip in Rrhs1 was present in the *R. solanacearum* Molk2 strain isolated from banana in the Philippines [38, 70, 71]. Learning how, why, and when Molk2 acquired this Rhs-CT tip and if it contributes to a competitive fitness against other bacteria in banana production remains to be elucidated.

The Rrhs2 tips are predicted DNases encoded by a GH-E motif and were present in 34 strains of *R. pseudosolanacearum,* making it the second most abundant tip for the genus [10, 24] (Fig 4B, Fig S29; Table S3). Like Rrhs2 tips, Rrhs5, Rrhs10, and Rrhs11 were conserved in only specific species (Table S3). The Rrhs5 tips are only present in *R. solanacearum*, while the tips in Rrhs10 and Rrhs11 were only present in *R. syzygii* (Table S3). The third most abundant Rhs-Ct tip in the RSSC was Rrhs3, present in 16 strains, including 14 *R. pseudosolanacearum* and two *R. syzygii* strains (Fig. 3B). This cluster, like Rrhs2, Rrhs4, Rrhs8, Rrhs12, Rrhs20, and Rrhs27, are predicted DNases [11, 24] (Fig 4B; Fig S29; Table S3).

A total of 106 unique Rhs-CT tips, named Prhs1-Prhs106, were identified among the 255 Rhs-CT tip sequences retrieved from the 60 *Pectobacterium* genomes. Of those, 42 were singletons present in only one strain of the genus (Fig 3C; Table S3). The remaining *Pectobacterium* Rhs-CT tips were present in at least two genomes. Specifically, 34 Rhs-CT tips were present in two *Pectobacterium* strains, 27 Rhs-CT tips were shared among three to seven strains, and only three tips were share among eight or more strains (Fig 3C; Table S3). The three most predominant Rhs-CT tips among *Pectobacterium* pathogens were Prhs1, Prhs2, and Prhs3. Each of these clusters had eight Rhs-CT tips encoded in the genomes of different *Pectobacterium* species. The Prhs1 tips encode a putative toxin of unknown function. Furthermore, these eight Rhs-CT tips are only associated with full-length Rhs toxin sequences in the genomes of *P. versatile, P. carotovorum,* and *P. brasiliense* (Fig 4C; Fig S30; Table S3). The tips in Prhs2 encode a DNase containing a ParB motif. Six of the eight Prhs2 tips are associated with orphan module sequences encoded in the genomes of *P. parmentieri* (Fig 4C; Fig S30; Table S3). The remaining two tips are associated with pseudogene sequences and are encoded in the genome of *P. carotovorum* strain IPO:4062 NAK:237 and *P. brasiliense* strain WPP14. Like Prhs2, tips in Prhs3 encode a DNase and were extracted from genomes of *P. parmentieri*, *P. versatile*, *P. carotovorum*, and *P. aroidearum* (Fig 4C; Fig S30; Table S3). Five of eight Prhs3 tips are associated with orphan module sequences encoded in the genomes of *P. parmentieri* and *P. aroidearum*, two are associated with full-length Rhs toxin sequences in the genomes of *P. versatile* and *P. carotovorum,* and one tip is associated with a predicted pseudogene sequence in *P. versatile* (Fig 4C; Fig S30; Table S3).

A total of 46 unique Rhs-CT tips, named Drhs1-Drhs46, were identified among the 113 Rhs-CT tip sequences retrieved from 31 *Dickeya* genomes. Of these, 16 were singletons present in only one *Dickeya* strain (Fig 3D; Table S3). The remaining *Dickeya* tips were present in at least two strains of the genus. Specifically, 13 Rhs-CT tips are present in two *Dickeya* strains, 16 tips are shared among three to seven strains, and only one tip was shared among eight strains (Fig 3D; Table S3). *Dickeya* had the lowest number of predicted Rhs CT tips (Fig 1A; Table S3). This finding is likely an artifact of the fewer publicly available closed genomes for *Dickeya* in NCBI (Table S1 and S2). Additional genomes from isolates collected in different locations and hosts will help confirm whether *Dickeya* strains have lower Rhs toxin abundance and functional diversity. This finding would suggest that *Dickeya* strains are under lower competitive pressures in their microbe-rich environments.

Drhs1 was the most abundant Rhs-CT tip among *Dickeya* strains. It was encoded in *D. dadantii*, *D. dianthicola, D. fangzhongdai*, and *D. solani.* Interestingly, *D. dadantii* strain 3937 had two Rhs-CT tips in this cluster, one associated with a full-length Rhs toxin sequence and the other with an orphan module (Fig 4D; Fig S31; Table S3). This phenomenon, where the same Rhs-CT tip was found to be associated with both a full-length Rhs toxin sequence and a separate orphan module in the same genome, was observed in *P. parmentieri* (HC and IFB548), *P. aroidearum* (AK042), *P. atrosepticum* (Green1 and SCRI1043), *P. carotovorum* (25.1), *X. vesicola* (NCPPB 2649 and SV1601), and *X. oryzae* (BB151-3) (Fig 4A and 4D). A total of 16 clusters in this genus are predicted to encode DNases via different toxin domains, such as AHH, HNHc, WHH, Tox-SHH, Tox-HNH-EHHH, ParB_N._Srx, Colicin-DNase, GIY-YIG_SF, NUC and LHH [10] (Fig 4D; Fig S31; Table S3). Like all other genera in this study, the predicted function of most Rhs-CT tips in *Dickeya* is unknown (Fig 4; Table S3).

### Most Rhs-CT tips in *Xanthomonas*, *Ralstonia*, *Pectobacterium*, and *Dickeya* remain uncharacterized, highlighting a significant knowledge gap

The expected function of 69.64% of all C-terminal tip sequences across all genera remains unknown. This includes 461 individual Rhs-CT tip sequences across 147 unique Xrhs tip clusters, 191 Rhs-CT tip sequences across 58 unique Rrhs tip clusters, 151 Rhs-CT tip sequences across 68 Prhs tip clusters, and 47 Rhs-CT tip sequences across 20 Drhs tip clusters (Fig 4; Table S3). DNases were the second most prominent enzymatic activity encoded by Rhs-CT tips (Fig 4). This includes 79 individual Rhs-CT tip sequences across 18 Xrhs tip clusters in *Xanthomonas*, 89 individual Rhs-CT tip sequences across 13 Rrhs tip clusters in *Ralstonia*, 71 individual Rhs-CT tip sequences across 26 Prhs tip clusters in *Pectobacterium*, and 60 individual Rhs-CT tip sequences across 24 Drhs tip clusters in *Dickeya*. RNases were the third most prominent enzymatic activity encoded by Rhs-CT tips across all genera (Fig 4; Table S3). However, this enzymatic activity was significantly lower than Rhs-CT tips predicted to be DNases and those with unknown function. Predicted RNases include 23 individual Rhs-CT tip sequences across five Xrhs tip clusters in *Xanthomonas*, eight individual Rhs-CT tip sequences across four Rrhs tip clusters in *Ralstonia*, 22 individual Rhs-CT tip sequences across eight Prhs tip clusters in *Pectobacterium*, and six individual Rhs-CT tip sequences across two Drhs tip clusters in *Dickeya* (Table S3).

### Proteases and deaminases are notably underrepresented among Rhs-CT tips in the genomes of *Xanthomonas*, *Ralstonia*, *Pectobacterium*, and *Dickeya*

The least common enzymatic function predicted for Rhs-CT tips across the four studied genera are proteases and deaminases (Fig 4). A total of 15, six, and seven individual protease-encoding Rhs-CT tip sequences grouped into 11, one, and four clusters for *Xanthomonas, Ralstonia,* and *Pectobacterium*, respectively (Fig 4; Table S3). One protease encoding Rhs-CT tip was found in one genome of *X. euroxanthea, X. citri, X. arboricola, X. theicola, X. albilineans, X. oryzae,* and *X. fragariae,* and two genomes of an unclassified *Xanthomonas* species (Table S3).

Notably, two distinct protease encoding Rhs-CT tips were found in the genome of *X. axonopodis* pv. *vasculorum* NCPPB 796, the causal agent of gumming disease in sugarcane and isolated in Brazil (Table S2 and S3). In this genome, one Rhs-CT tip was associated with a full-length Rhs toxin sequence and the other with a pseudogene. The remaining three protease-encoding Rhs-CT tips were found in three strains of *X. sacchari* isolated from rice microbiomes. Two of these strains, HR3-46 and HR1-32, were isolated in Yunnan province, and one, YT9-19-2, in SanYa province, both in Southern China (Table S2 and S3).

Interestingly, *X. sacchari* strains isolated from rice in Northern China did not contain protease-encoding Rhs-CT tips in their genomes. This finding indicates that Rhs toxins contribute to *Xanthomonas* niche adaptation.

One protease Rhs-CT tip cluster, Rhs11, was found exclusively in *R. syzygii* genomes and contained a Peptidase_C80 toxin domain (Fig 4B; Fig S29; Table S3). Curiously, Rhs11 tip sequences were only associated with pseudogenes (Table S3). In *Pectobacterium,* the protease Rhs-CT tips were encoded in the genomes of *P. versatile*, *P. brasileiense*, *P. aroicearum*, and *P. parmentieri* either as a full-length Rhs toxin, pseudogene, or orphan module (Table S3).

Notably, only one of the numerous strains from each of these species had a single protease encoding Rhs-CT tip, except for *P. parmentieri* strain HC, isolated from a potato in South Korea, that carried both a full-length Rhs toxin and an orphan module associated with protease activity (Table S2 and S3). No protease Rhs-CT tips were identified in *Dickeya* genomes in this study. However, the presence of 20 Rhs-CT tip clusters with unknown functions leaves open the possibility that one may encode a yet-undiscovered protease toxin domain.

Rhs-CT tips with deaminase-predicted functionality were only identified in *Xanthomonas* and *Pectobacterium* genomes (Fig 4; Table S3). In *Xanthomonas,* the 26 deaminase Rhs-CT tips clustered into Xrhs4 and Xrhs12 and were extracted from the genomes of *X. oryzae* and *X. translucens,* both pathogens of monocotyledon plant hosts (Table S3). All 26 Rhs-CT tips in these two tip clusters are associated with only full-length Rhs toxin sequences (Fig 4; Table S3). The only other deaminase-predicted Rhs-CT tip cluster was Prhs 24, identified in *Pectobacterium*. This cluster contained four Rhs-CT tip sequences featuring a YwqJ- toxin domain, previously described by Iyer et al. (2011) in *Bacillus,* a Gram-positive soil-dwelling bacterium (Table S3). This finding supports the hypothesis that Rhs-CT tips are exchanged among members of microbial communities through horizontal gene transfer. Two of the four Rhs-CT tip sequences in Prhs24 are associated with full-length Rhs-toxin genes in *P. versatile* 45353 and *P. carotovorum* PC1 genomes. The remaining two tips are linked to pseudogenes in the genomes of *P. versatile* F131 and SR1 (Table S3).

### Select Rhs-CT tips are exclusively associated with either full-length Rhs toxin, pseudogene, or orphan module sequences, underscoring their functional and evolutionary specificity

Our analysis identified Rhs-CT tips associated exclusively with either full-length Rhs-toxin genes, pseudogenes, or orphan module sequences within specific bacterial genera. For instance, 39, 13, 27, and 13 Rhs-CT tip clusters are linked solely to full-length Rhs-toxin sequences in the genomes of *Xanthomonas*, *Ralstonia*, *Pectobacterium*, and *Dickeya*, respectively (Fig 4; Table S3). These full-length Rhs toxins have the genetic elements necessary for their immediate translocation into a target host cell, indicating their enzymatic actions are crucial for the competitive fitness of these strains within their current microbiomes.

Similarly, one, three, and 18 Rhs-CT tips are associated exclusively with pseudogenes in *Xanthomonas*, *Ralstonia*, and *Pectobacterium*, respectively (Fig 4; Table S3). This finding suggests that the enzymatic functions of these toxins no longer provide a competitive advantage and are gradually being eliminated from these strains’ genomes in their current microbial environments. Additionally, ten, two, and six Rhs-CT tip clusters are associated with only orphan module sequences in *Xanthomonas*, *Ralstonia*, and *Pectobacterium*, respectively (Fig 4; Table S3). Orphan modules are likely maintained because they provide **enzymatic diversity** and **metabolic flexibility**, which can be critical for adapting to fluctuating environments or for capitalizing on new ecological opportunities. These modules might not be actively used all the time but could become valuable if environmental conditions change or if the strain encounters new competitors, substrates, or stressors. This diversity allows bacteria to remain versatile, which is important for survival in complex, competitive microbial communities. Notably, this analysis found no Rhs-CT tip clusters exclusively associated with pseudogenes or orphan module sequences in *Dickeya* genomes.

Not surprisingly, we also found instances where the same Rhs-CT tips was associated with a full-length Rhs toxins sequence in one strain, a pseudogene in a second strain, and/or to an orphan module in a third strain from either the same or different species (Fig 4;Table S3). This was observed in *Xanthomonas* Rhs-CT tips Xrhs1, Xrhs3, and Xrhs13; *Ralstoina* Rhs-CT tips Rrhs6, Rrhs9, and Rrhs26; and *Pectobacterium* Rhs-CT tips Prhs2-Prhs7, and Prhs9 (Table S3). These data support the hypothesis that Rhs-CT tip displacement is a natural phenomenon in bacterial genomes and that competitive pressures most likely determine Rhs-CT tips genomic location within Rhs toxin systems [10]. We also demonstrate that Rhs-CT tips can be shared among phytopathogenic bacterial strains both intra- and interspecifically (Fig 4E; S32; Table S3).

### Select Rhs-CT tips are shared across bacterial genera

To investigate the potential for Rhs-CT tip exchange at the intergenus level, we performed an all-vs-all BLASTP analysis of all 1,267 Rhs-CT tip sequences from the four genera. We identified five clusters of Rhs-CT tips that shared over 50% identity between *Ralstonia* and *Xanthomonas* strains (Fig 4E; S32; Table S3). One of the clusters with unknown function included eight Rhs-CT tips: five from *Xanthomonas* Xrhs43 and three from *Ralstoina* Rrhs22. These were extracted from *X. oryzae* and *R. pseudosolanacearum* strains (Table S3). The next two clusters included two Rhs-CT tips from *Xanthomonas* and two from *Ralstonia* (Fig 4E; S32; Table S3). Intriguingly, in each cluster, the two *Xanthomonas* Rhs-CT tips were extracted from the genomes of the same *X. sacchari* strains (HR1-32 and YT9-19-2) isolated from rice China (Table S3). While the two *Ralstonia* Rhs-CT tips were extracted from the genomes of *R. solanacearum* strain RUN2474 and *R. syzygii* strain LLRS-1, isolated from Madagascar and China for one cluster; and *R. pseudosolanacearum* strain RUN2340 and *R. solanacearum* CFBP2957 isolated in Madagascar and the West Indies for the second cluster (Table S3).

One cluster sharing more than 60% identity was found among Rhs-CT tips from *Ralstonia* and *Pectobacterium*. This cluster has nine Rhs-CT tips, five from *Ralstoina* Rrhs28 and four from *Pectobacterium* PRhs16 (Fig 4E; S32; Table S3). The Rhs-CT tips in this cluster were extracted from only *P. parmentieri* genomes. Two of the four Rhs-CT tips were associated with orphan module and two with pseudogenes. The *Ralstonia* Rhs-CT tips include four extracted from *R. solanacearum* genomes associated with full-length Rhs toxin sequences only. The fifth *Ralstonia* tip was extracted from *R. syzygii* and also associated with a full-length Rhs-toxin sequence (Table S3).

One cluster sharing more than 90% identity was found among Rhs-CT tips from *Ralstonia* and *Dickeya* (Fig 4E; S32; Table S3). This cluster had four Rhs-CT tips, two from *Ralstonia* Rrhs37 and two from *Dickeya* Drhs25, all are predicted DNases and associated with full-length Rhs toxin sequences. The *Ralstonia* Rhs-CT tips in Rrhs37 were extracted from *R. solanacearum* strains Rs5 and K60, isolated in the U.S. from tomato plants. These two genomes are the only available representatives for the pathogenic *Ralstonia* population endemic to the Southeastern U.S. The two *Dickeya* Rhs-CT tips were extracted from the genomes of *D. dadantii* strain M2- 3, isolated from potato in China, and *D. dianthicola* strain LAR.16.03.LID, also isolated from potatoes in Morocco (Table S2).

Finally, we identified 19 clusters containing Rhs-CT tips from *Pectobacterium* and *Dickeya* that share over 90% identity (Fig 4E; S32; Table S3). The largest cluster contains 14 Rhs-CT tips, six from *Pectobacteirum* Prhs7 and eight from *Dickeya* Drhs1, all encode a DNase with a NUC toxin domain. Tips in this cluster were associated with full-length Rhs toxin genes, pseudogenes, and orphan module sequences (Table S3). Among the Rhs-CT tips in this cluster, two alleles of one Rhs-CT tip were identified in *D. dadantii* 3937. One allele is associated with a full-length Rhs toxin sequence RhsA (DDA3937_RS04095), while the other is associated with an orphan module (DDA3937_RS23790) not previously detected in this strain’s genome (Koskiniemi et al. 2013b) (Table S3).The two additional full-length Rhs toxins previously reported for this strain were also retrieved with our Rhs HMM model, RhsB (DDA3937_RS13755) and RhsC (DDA3937_RS06980), and the two orphan modules rhs_Co1_ (DDA3937_RS06990) and Rhs_Co2_ (DDA3937_RS06995) [9]. RhsA and RhsB are confirmed DNases encoded by a NUC and HnHc toxin domain. The function of RhsC remains unknown.

Interestingly, despite being isolated from different plant hosts and geographic location, RhsC was also found in the genome of *D. dadantii* strain XJ12, the causal agent of bacterial sheath rot of banana [72]. Like *D. dadantii* 3937, XJ12 also carries the two orphan modules associated with RhsC: *rhsC_o1_* and *rhsC_o2_* [9]. The *rhsC_o1_* is also present in the genomes of *D. zeae* EC2 and *D. fangzhongdai* ZXC1; however, it is a full-length Rhs toxin gene and not an orphan module in these strains. In *D. zeae* MS-2018 and *D. fangzhongdai* Onc5, rhs_Co1_ is associated with pseudogene sequences. The rhs_Co2_ was also extracted from the genomes of *D. chrysanthemi* Ech1591 as a full-length Rhs toxin gene and in *D. zeae* EC1 as an orphan module (Table S3).

## Conclusion

Using an Rhs-HMM model, we analyzed the abundance and diversity of Rhs polymorphic toxins across four genera of plant pathogenic bacteria—*Xanthomonas*, *Ralstonia*, *Pectobacterium*, and *Dickeya*—using publicly available genomes. Our data suggest that Rhs toxins are secreted via more than one secretion system, including the T2SS, and not only the canonical T6SS. We found no clear association between Rhs toxin profiles and factors such as geographic origin or pathogen lifestyle. Interestingly, *Xanthomonas* displayed the highest Rhs toxin abundance, contradicting our initial hypothesis that soilborne pathogens like *Ralstonia* would carry more diverse and abundant Rhs toxins, given the high bacterial density in soil (estimated at 10¹⁰ bacteria per gram) [73]. Expanding this analysis to include more genera of foliar plant pathogens may help clarify these patterns. Additionally, our results may be influenced by biases in the availability and completeness of whole-genome sequences across these genera.

Our findings corroborate previous research on the widespread presence of Rhs toxins across bacterial genera. Notably, we also identify certain species of plant pathogenic bacteria that lack detectable Rhs toxin sequences. We confirmed the presence of homologous Rhs toxin sequences across multiple strains, species, and genera, reinforcing the hypothesis that these toxins are disseminated via horizontal gene transfer. Additionally, our data suggest that interactions within microbial communities may contribute to the diversification of Rhs toxins, even among distantly related strains. For instance, *Ralstonia* strains, rich in pseudogenes, suggest intense intraspecies rivalry, while *Pectobacterium* and *Dickeya* share toxin domains and immunity proteins, enabling coexistence. *Xanthomonas*’s unique singletons and diverse toxin functions (e.g., RNases, DNases) underscores the system’s hypervariability.

Insights into the competitive dynamics of plant pathogenic bacteria within complex microbial communities can improve understanding of pathogen survival and virulence in microbe-rich environments. By characterizing the abundance, diversity, and functions of Rhs toxin, we are beginning to answer the why and how pathogens deploy enzymatic proteins to eliminate microbial rivals, modulate their microbiomes, and successfully colonize plant hosts. These findings may reframe plant pathogens not as passive invaders but as active, strategic competitors engaging in an evolutionary arms race shaped by horizontal gene transfer, selective pressure, and niche specialization, repositioning disease emergence as a community-level phenomenon.

## Materials and Methods

### Data sources

Research has shown that in draft bacterial genomes, *rhs* toxin genes are often at the end of contigs, thus confounding the accuracy of predicted Rhs toxins in a strain, structure, and function [40]. To bypass this limitation, only whole genome sequences with a single complete circular chromosome for *Xanthomonas, Ralstonia*, *Pectobacterium*, and *Dickeya* were downloaded on June 25, 2023, from the NCBI public genome database (http://www.ncbi.nlm.nih.gov) in GenBank format (Table S1). Additionally, two genomes from *Escherichia coli* K-12 and *P. aeruginosa* PAO1 were retrieved and used as outgroups in phylogenetic analysis.

### Rhs-Hidden Markov Model

After a comprehensive literature review on functional studies of *rhs* genes, multiple Rhs amino acid sequences were manually extracted from the NCBI protein database. A total of 54 Rhs amino acid sequences from 27 strains belonging to nine distinct bacterial genera (Fig S33; Table S5) were aligned in Geneious Prime version 2021 (www.geneious.com) using the Clustal Omega (v1.2.3) algorithm with default parameters. The conserved motif that demarcates the hypervariable C-terminal tip from the N-terminal domain, approximately 10 amino acids in length (PxxxxDPxGL) [2, 9] was extracted and realigned. The realigned sequences were used to build a Hidden Markov Model (HMM) using the *hmmbuild* command in HMMER v3.3 (http://hmmer.org). To confirm the model’s accuracy in extracting *rhs* genes, the model was used to confirm *rhs* gene presence in the same strain used to generate the model (Table S5) and three *R. solanacearum* species complex strains, K60, UW551, and GMI1000. This method was then repeated by adding amino acids to the 5’ end of the 10-aa motifs until an HMM was generated that accurately extracted all putative Rhs toxins from bacterial genomes. After manual curation and genome mining, a final Rhs HMM, from here on referred to as Rhs HMM, was built using a 43-aa peptide motif that included the original 10-aa motif and 33-aa upstream to it (Fig S33; Table S5).

### Mining whole-genome sequences (WGS) for *rhs* Genes

An average nucleotide identity (ANI) analysis was conducted on all downloaded genomes for each of the four bacterial genera. If any genome clustered into an ANI group with 100% similarity, they were considered clones, and all but one were removed from further study (Table S1). The remaining WGS were translated using a custom Perl script (https://github.ncsu.edu/Huerta-lab/Mine_RhsToxins/blob/main/gb2cds.pl). The Rhs HMM was then used to scan all genomes from each genera using the *hmmsearch* command of HMMER v3.3 using and E-value of 0.05. Returned sequences were passed through the NCBI Conserved Domains database to confirm the presence of RHS domains to then be manually curated [74]. Rhs toxins that lacked a full-length C-terminal tip (<90 aa), or were composed just by the motif (i.e., only the 43-aa sequence was retrieved) were removed from further study (Table S2).

### Rhs toxin families

To cluster Rhs toxins into families, we followed the methods described by Jackson et al [2]. Briefly, the Rhs sequences from each genus were aligned to identify and mark the 10-aa peptide motif that demarcates the conserved N-terminal from the hypervariable Rhs-CT tip. Any sequence before the motif, considered the Rhs N-terminal, was extracted and realigned using MAFFT v.7.511 [75] following the auto-alignment strategy implemented in the DeCIFR toolkit [76]. Alignments were visualized in Geneious Prime v10.1 (https://www.geneious.com) to identify and extract the Rhs “core region,” defined as the sequence flanked by the two amino acid motifs RvxxxxxxxG and PxxxxDPxGL [2]. The Rhs “core region” sequences within each genus were aligned using MAFFT v.7.526 and maximum likelihood phylogenies were inferred using both RAxML v8.2.12 [77], with a general time reversible (GTR) matrix-based model of amino acid substitution rates and 1,000 bootstrap replicates [50] and IQ-TREE [78] with model selection and bootstrapping implemented in the DeCIFR toolkit [76]. The phylogenies were visualized in the DeCIFR Tree-Based Alignment Selector (T-BAS, version 2.4) toolkit [79–81]. Resulting clades with >90% and >95% bootstrap support in RAxML and IQ-TREE, respectively, were used to define Rhs toxin families for each genus as described in Jackson et al. [2]. To compare and visualize Rhs toxin system clusters among and across bacterial species and genera, a custom python script was generated to extract the 5000 bp and 3000 bp region upstream and downstream of each Rhs “core region,” respectively (https://github.ncsu.edu/Huerta-lab/Mine_RhsToxins/blob/main/Extracting_Rhs_Down_Ups.py). For each family, the sequences were extracted and saved in FASTA format and uploaded to Clinker (v0.0.26) clustermap.js [82] to visualize synteny among Rhs toxin system loci in bacterial genomes.

### C-terminal tip functional diversity prediction

After Rhs sequences from each genus were aligned and the 10 aa motif was identified, any sequence after, including the 10-aa motif, was considered the Rhs-CT tip and was extracted and realigned as described above. This alignment was used to run a pairwise BlastP analysis to create a similarity network analysis (SNA) using the Enzyme Similarity Tool (EFI-EST) [83–85]. The SNA was saved in xgmml format and visualized in Cytoscape v3.3.0. [86] (Fig 4; Fig S28-S32). The Rhs-CT tips were then run on the NCBI Conserved search domain (CDD) and Pfam database to assign predicted functions to all Rhs-CT tip sequences and clusters in this study [74, 87] (Table S3).

## Acknowledgements

The authors thank Dr. David Ritchie for insightful discussion and manuscript feedback.

## Supporting Information Captions

**Table S1. Bacterial genomes used in this study.**

**Table S2. Genomes used in this study pre and post data curation.**

**Table S3 Rhs toxin sequence characteristics and enzymatic functions.**

**Table S4. Clade designations for Rhs toxin families.**

**Table S5. Strains and sequences used to generate Rhs HMM.**

**Fig S1.** IQ-TREE (A) and RaxML (B) phylogenies with clade support for Rhs toxin N-terminals in *Xanthomonas* strains. Bootstrap support values are displayed on the branches.

**Fig S2.** IQ-TREE (A) and RaxML (B) phylogenies with clade support for Rhs toxin N-terminals in *Ralstonia* strains. Bootstrap support values are displayed on the branches.

**Fig S3.** IQ-TREE (A) and RaxML (B) phylogenies with clade support for Rhs toxin N-terminals in *Pectobacterium* strains. Bootstrap support values are displayed on the branches.

**Fig S4.** IQ-TREE (A) and RaxML (B) phylogenies with clade support for Rhs toxin N-terminals in *Dickeya* strains. Bootstrap support values are displayed on the branches.

**Fig S5.** High-resolution image of Fig 2A.

**Fig S6.** High-resolution image of Fig 2C.

**Fig S7.** High-resolution image of Fig 2D.

**Fig S8.** High-resolution image of Fig 2E.

**Fig S9.** High-resolution image of Fig. 2B gene synteny among Rhs toxin loci found in XRhs **F I.**

**Fig S10.** High-resolution image of Fig. 2B gene synteny among Rhs toxin loci found in XRhs F II.

**Fig S11.** High-resolution image of Fig. 2B gene synteny among Rhs toxin loci found in XRhs F III.

**Fig S12.** High-resolution image of Fig. 2B gene synteny among Rhs toxin loci found in XRhs F IV.

**Fig S13.** High-resolution image of Fig. 2B gene synteny among Rhs toxin loci found in XRhs F V.

**Fig S14.** High-resolution image of Fig. 2B gene synteny among Rhs toxin loci found in XRhs F VI.

**Fig S15.** High-resolution image of Fig. 2B gene synteny among Rhs toxin loci found in XRhs F VII.

**Fig S16.** High-resolution image of Fig. 2B gene synteny among Rhs toxin loci found in XRhs F VIII.

**Fig S17.** High-resolution image of Fig. 2F gene synteny among Rhs toxin loci found in RRhs F I.

**Fig S18.** High-resolution image of Fig. 2F gene synteny among Rhs toxin loci found in RRhs F II.

**Fig S19.** High-resolution image of Fig. 2F gene synteny among Rhs toxin loci found in RRhs F III.

**Fig S20.** High-resolution image of Fig. 2G gene synteny among Rhs toxin loci found in PRhs F I.

**Fig S21.** High-resolution image of Fig. 2G gene synteny among Rhs toxin loci found in PRhs F II.

**Fig S22.** High-resolution image of Fig. 2G gene synteny among Rhs toxin loci found in PRhs F III.

**Fig S23.** High-resolution image of Fig. 2G gene synteny among Rhs toxin loci found in PRhs F IV.

**Fig S24.** High-resolution image of Fig. 2G gene synteny among Rhs toxin loci found in PRhs F V.

**Fig S25.** High-resolution image of Fig. 2H gene synteny among Rhs toxin loci found in DRhs F I.

**Fig S26.** High-resolution image of Fig. 2H gene synteny among Rhs toxin loci found in DRhs F II.

**Fig S27.** High-resolution image of Fig. 2H gene synteny among Rhs toxin loci found in DRhs F III.

**Fig S28.** High-resolution image of Fig 4A.

**Fig S29.** High-resolution image of Fig 4B.

**Fig S30.** High-resolution image of Fig 4C.

**Fig S31.** High-resolution image of Fig 4D.

**Fig S32.** High-resolution image of Fig 4E.

**Fig S33.** Schematic representation of methods used to generate the Rhs HMM. (A) A 43-aa-long sequence was extracted from 54 strains across nine bacterial genera. These sequences were aligned to build an Rhs HMM using the hmmbuild command in HMMER v3.3. (B) Weblogo image depicting conserved residues derived from the multiple sequence alignment of 54 representative Rhs toxin homologs. The x-axis represents the positions in the motif, and the y-axis reports the conservation in bits.

